# Maladaptive coping strategies and neuroticism mediate the relationship between 5HTT-LPR polymorphisms and symptoms of anxiety in elite athletes

**DOI:** 10.1101/493320

**Authors:** Mario Altamura, Salvatore Iuso, Giovanna D’Andrea, Francesca D’Urso, Carla Piccininni, Eleonora Angelini, Francesco Sessa, Maurizio Margaglione, Caterina Padulo, Beth Fairfield, Annamaria Petito, Antonello Bellomo

## Abstract

Previous studies have suggested that genetic factors, personality traits and coping strategies might play independent and interacting roles in influencing stress-related anxiety symptoms. The aim of this study was to examine whether Neuroticism and maladaptive coping strategies mediate the association between the serotonin transporter gene-linked polymorphic region (5HTT-LPR) and symptoms of anxiety and depression in elite athletes who experience high levels of competitive stress. One hundred and thirty-three participants were genotyped for the 5-HTTLPR polymorphism and then asked to complete the Cope Orientation to Problems Experienced Inventory and the NEO Five-Factor Inventory. A path analysis was used to test the aforementioned hypothesis. The 5HTT-LPR was significantly associated with Neuroticism, the coping strategy of Focus on and Venting of Emotions’ (FVE) and symptoms of anxiety. FVE and Neuroticism mediated the association between the 5HTT-LPR and symptoms of anxiety (i.e., Cognitive Anxiety and Emotional Arousal Control). Also, Neuroticism was a mediator of the association between the 5HTT-LPR and FVE. Finally, FVE also mediated effects on the relationship between Neuroticism and symptoms of anxiety. Results suggest that the 5HTT-LPR may affect the susceptibility to develop symptoms of anxiety in elite athletes indirectly through mediation by FVE and Neuroticism.

## Introduction

A growing body of evidence suggests that elite athletes may suffer from subsyndromal symptoms of depression and anxiety (i.e., symptoms not meeting the threshold for a formal DSM diagnosis) as a result of significant sport-related stressors (e.g., injury, performance failure, over training, club/organizational climate, negative group dynamics/peer interaction) [1, 2, 3, 4, 5, 6]. Indeed, cross-sectional [7, 8, 9, 10, 11]; and prospective cohort studies [12, 13] (among elite athletes show a prevalence of anxiety/depression symptoms (quantified on a rating scale) in the range of 15 to 57%. Explanations for this prevalence comprise a number of barriers including stigma and a lack of awareness [14, 15]. Although there is evidence that elite athletes are highly vulnerable for developing mental health problems due the unique range of stressors involved [16, 17], less is known about the potential determinants and mechanisms linking sport related stress to symptoms of depression and anxiety in this population. More importantly, since the development of anxiety/depression symptoms can be described by a multifactorial model of causality, biological (e.g. genetics), psychological (e.g. personality traits, coping strategies) and social (e.g., social supports) factors may all interact to contribute to the development of symptoms of anxiety and depression in elite athletes [18, 19, 20, 21].

The serotonin transporter protein (5-HTT), which controls the reuptake of serotonin (5-HT) from the synaptic clefts, plays a critical role in regulating many physiological functions including cognition and emotional behavior [22, 23]. In humans, the transcriptional activity of the gene encoding 5-HTT is modulated by a polymorphism in the promoter region of the gene, designated 5HTT-LPR [24]. Specifically, the short (s) allele (5HTT-LPR-s) is associated with lower transcriptional efficiency of 5-HTT compared to the long (l) allele. This results in a less efficient transporter function and, consequently, in a net reduction in circulating serotonin [25, 26]. The 5HTT-LPR genotype has been widely studied and many studies have reported associations between the 5HTT-LPR genotype and anxiety-related personality traits (i.e., Neuroticism) and risk for multiple affective disorders, including symptoms of anxiety and depression [27, 28, 29]. Moreover, this association may be even stronger in the context of stressful life events [30, 31] Other studies, instead, have reported a more inconsistent association, revealing how the exact nature of the role of 5HTT-LPR genotype in abnormal mood and anxiety remains somewhat unclear [see for meta-analysis, 32]. Noteworthy, recent studies reported that 5-HTTLPR differences in anxiety and depression symptoms may be best observed in the context of everyday environmental stressors [33, 34]. In line with this biological “stress reactivity”, several studies have found that, individuals homozygous for the s allele exhibit greater fMRI amygdale activation compared to individuals with a single allele [35, 36] and HPA-axis reactivity [37] in response to immediate threat, has been confirmed in mouse models [38]. Therefore, based on these findings we expect genotype differences in anxiety and depression symptoms to emerge significantly in the context of acute stress in competitive sport. Another possible explanation for the mixed results is that genetic vulnerability may influence the development of symptoms of anxiety and depression only when entirely or partially mediated by psychosocial factors (e.g., trait coping style, personality) [39, 40].

Coping reflects conscious attempts to manage stress in the context of a situation or condition perceived as challenging, or that exceeds an individual’s resources to adequately deal with a problem [e.g., 41]. Coping strategies can be conceptualized as “problem focused” (i.e., attempting to control the problem causing the distress by modifying oneself or ones environment) or “emotion focused” (i.e., utilizing coping strategies to reduce the negative emotional responses associated with stress) [42]. Problem-focused coping is generally viewed as an adaptive mode of coping that involves actively planning or engaging in a specific behavior to overcome the problem causing distress. Emotion-focused coping strategies, instead, are often divided into two sub categories; active emotion-focused coping and avoidant emotion-focused coping. Active emotion-focused (e.g., positive reframing) is generally viewed as being an adaptive emotion regulation strategy whereas avoidant emotion-focused coping (e.g., self-distraction) is seen to be maladaptive [43]. In addition, some studies have suggested that the use of emotion-focused coping styles in response to some types of stressors may be a risk factor for the development of affective disturbances [44]. Indeed, maladaptive emotion-focused strategies such as venting, denial, substance use, behavioral disengagement, self-distraction, self-blame have been shown to be associated with higher levels of anxiety and depression in both nonclinical [45, 46] and clinical samples [47, 48, 49]. In particular, sport related research on coping has frequently reported that elite athletes employ both adaptive problem-focused (e.g., problem solving, planning) and active emotion-focused coping (e.g., seeking emotional social support) strategies [50, 51, 52, 53] to cope with various sport-related stressors [see for reviews, 6, 54]. However, elite athletes also showed a tendency to use more avoidant coping strategies in training situations and during competition when they appraised the stressor as a threat [55] or when faced with unexpected stressors [56, 57]. Moreover, several studies demonstrated that competitive athletes who report more maladaptive emotion-based coping strategies tend to experience greater overuse injury risk [58] and symptoms of anxiety and depression [7, 59, 60]. Perhaps most notably, recent studies suggest that the 5-HTT-LPR s*-*allele could strongly influence the differentiation of individual trait coping styles [40, 61, 62]. Specifically, these studies reported that 5HTT-LPR-s allele carriers demonstrated lower scores in positive coping styles (i.e., problem solving strategies) and higher scores in maladaptive trait coping styles than ll genotype individuals. Furthermore, Wang et al. [40] reported that maladaptive coping mediates the association between the 5HTT-LPR-s and lifetime major depression in adults. Therefore, it is conceivable to hypothesize that maladaptive coping styles may entirely, or partially, mediate associations between the 5HTT-LPR-s allele and anxiety/depression symptoms.

In addition to the inability to effectively cope with emotional experiences, studies have shown that negative personality traits such as Neuroticism might influence the development of symptoms of various clinical disorders including depression and anxiety disorders in response to recent stress [see for review, 63]. Moreover, prior genetic studies have shown associations between neuroticism and the s allele of the 5-HTT-LPR [64, 65, 66]. Furthermore, and most importantly, a number of studies have suggested that Neuroticism may mediate association between the 5HTT-LPR genetic variance and the development of depression [39, 40]. Over the past decades research on personality psychology has shown that Neuroticism and dysfunctional coping strategies might play both independent and interacting roles in influencing psychophysiological responses to stressful experiences and the development of mental health problems [see for review, 67]. In particular, a recent meta-analysis reported that maladaptive emotion-focused coping (i.e., the expression of negative emotions) related positively (and strongly) to Neuroticism [68]. There is also evidence that the relation between Neuroticism and anxiety/depression phenotypes is mediated by individual differences in the use of different emotion regulation strategies. Specifically, it has been suggested that the use of maladaptive coping mediated the association between Neuroticism and symptoms of depression [69] and emotional outcomes in a potentially stressful situation [70]. Together, these findings suggest that, besides the genetic influence, Neuroticism and maladaptive coping styles may provide significant and independent contributions to susceptibility to anxiety/depression phenotypes and mediate the association between 5HTT-LPR genotype and symptoms of anxiety and depression, which is consistent with the complexity of these multi-factorial conditions.

The purpose of this study is to determine the mediating effects of maladaptive coping (i.e., emotion-focused coping strategies) and anxiety-related personality traits (i.e., Neuroticism) on the relationship between serotonin transporter gene promoter polymorphisms and anxiety/depression symptoms in elite athletes engaged in competitive sports. Our initial study on elite athletes suggested a direct association between 5HTT-LPR-s and the development of anxiety symptoms during competition; the short allele was related to anxiety symptoms in an additive fashion. Levels of anxiety symptoms were lowest among individuals with a ‘l/l’ genotype, intermediate with a s/l genotype, and highest with a s/s genotype. Furthermore, we demonstrated that Neuroticism (i.e. elevated neuroticism in s/s individuals) mediated the association between the 5HTT-LPR-s genotype and anxiety symptoms [19]. In this study, in addition to Neuroticism, we also investigated the possible mediation effects of maladaptive coping strategies in contribution of 5HTT-LPR polymorphisms in the development of symptoms of anxiety and depression (Fig. 1). Given existing evidence for an association between 5HTT-LPR genotype and maladaptive coping [40, 61] it is plausible to hypothesize that maladaptive coping might mediate the association between the 5HTT-LPR-s genotype and symptoms of anxiety and depression. Moreover, based on previous findings [e.g., 69] we hypothesized that maladaptive coping might also mediate the well-documented association between Neuroticism and symptoms of anxiety and depression. Finally, we investigated whether neuroticism mediates the association between the 5HTT-LPR genotype and maladaptive coping, but no prediction was made on that specific topic. A mediation analysis was used to test these hypotheses.

**Fig. 1.**
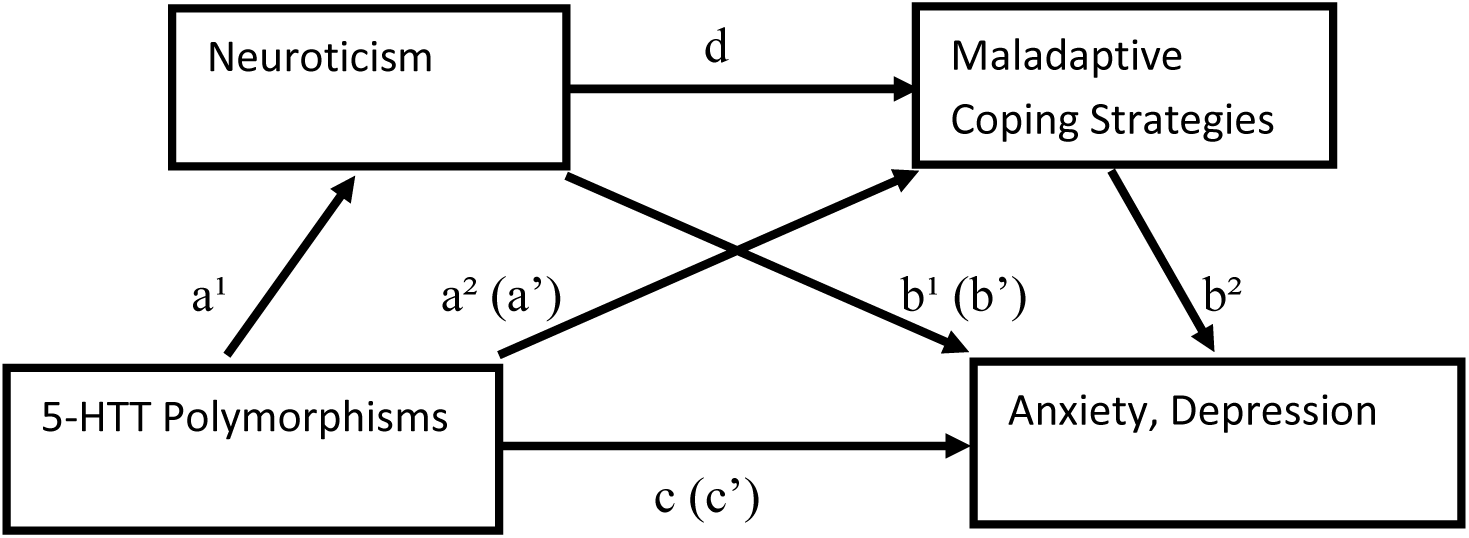
Path diagram of the multiple-mediation model for the study of 5-HTTLPR polymorphism, neuroticism, maladaptive coping styles, and symptoms of depression and anxiety.

## Methods

Participants in this sample were 133 elite male athletes between the ages of 18 and 36 (M = 23.36 (SD = 8.48 years). Recruitment and testing methods, fully reported in a previous study, are summarized as follows [19]. A letter of invitation to participate in the research project was sent to head coaches as well as to directors of elite sport organizations of the Apulia Region (Italy). Eighteen centers and 73% (133/182) of the elite male athletes agreed to participate in the study. Participants were athletes who competed at a national or international level in their chosen sport (i.e., soccer, basketball, hockey) and had more than two years of experience with national teams. All participants gave their written consent for taking part in the study, for collecting blood samples, as well as storing and subjecting it to a genetic analysis. Exclusion criteria included a history of traumatic brain injury, epilepsy, developmental disorder, diagnosable current substance abuse dependence or other known neurological condition and the presence or previous presence of psychiatric disorders. The Structured Clinical Interview for DSM-IV Axis I Disorders (SCID-I) [71] and Structured Clinical Interview for DSM-IV Axis II Personality Disorders (SCID-II) [72], were administrated to assess current and previous psychiatric diagnoses. The NEO Five-Factor Inventory (NEO-FFI) was used to assess personality [73]. Coping styles were measured using the Cope Orientation to Problem Experienced (COPE) inventory, a 60-item self-report designed to examine various coping styles used in response to stressful events [74]. For each item, response options are rated on 4-point scales from “never” to “always”. The inventory consists of 15 conceptually independent subscales, and provides scores on three main clusters of strategies (problem-focused coping; emotion-focused coping; maladaptive coping). Each cluster contains five specific scales. Five scales measure conceptually distinct aspects of problem-focused coping (active coping, planning, suppression of competing activities, restraint coping, seeking of instrumental social support); five scales measure aspects of active emotional-focused coping (seeking of emotional social support, positive reinterpretation and growth, acceptance, humor, turning to religion). Finally, five scales assess aspects of avoidant or maladaptive emotion-focus coping (focus on and venting of emotions, denial, behavioral disengagement, mental disengagement, alcohol/drug use). The Sport Performance Psychological Inventory (IPPS-48) and the Profile of Mood States (POMS) self-reporting questionnaires were used to assess anxiety and depressive symptoms. The IPPS-48 includes 48 items pertaining to eight factors. These factors are further divided into two broader conceptual categories. The Cognitive aspects category encompasses race preparation (RP), goal-setting (GS), mental practice (MP), and self-talk (ST) factors. The Emotional aspects category comprises self-confidence (SC), emotional arousal control (EAC), cognitive anxiety (CA) or worry and concentration disruption (CD). Low scores on one or more subscales reflect low-level mental skills, except for the subscales of CD, where high scores reflect difficulties. The IPPS-48 possesses a clear factorial structure, and good test-retest reliability [75]. The POMS is a standard validated psychological test consisting of 65 items that fit into 6 categories: tension-anxiety (T/A), depression-dejection (D/D), anger/hostility (A/H), vigor-activity (V/A), confusion-bewilderment (C/B) and fatigue-inertia (F/I) [76, 77]. All participants were also interviewed about significant sport-related stressors and feeling of stress in their own terms [78]. These unstructured interviews began with an open-ended prompt designed to guide the direction of the interview: “Tell me about your experience of stress as an athlete”. All of the athletes were participating in competitive sport events when the study was conducted. The present study is in accordance with the principles of the Declaration of Helsinki and was approved by the local Institutional Review Board (Comitato Etico ASL-FG; prot. n. 09/CE/07).

### DNA analysis

A blood sample was collected in ethylenediaminetetraacetic acid or sodium citrate from each participant and DNA was extracted from peripheral blood leukocytes according to standard protocols [79]. DNA amplification was amplified using the 2 flanking primers suggested in1996 by Heils et coll: 5-HTTU:5’GGCGTTGCCGCTCTUAATGC3’, nt-1416,-1397 5-HTTL:5’GAGGGACTGAGCTGGACAACCAC, nt-910,-889. This set of primers amplifies a 484/528 fragment corresponding to the SLC6A4_C short and long allele, respectively. The PCR reaction was carried out in a total volume of 20 μL consisting of 100 ng of genomic DNA, 0.1 μmol of primers per liter, 40-μmol/L deoxynucleotide triphosphates, 20-μmol/L 7-deaza-2’deoxyguanosine, and 1 unit of AmpliTaq with the appropriate buffer in a Mastercycler polymerase chain reaction thermal cycler (Eppendorf, Hamburg, Germany) [24]. Cycling conditions were as follows: 1 denaturing cycle at 95°C for 5 minutes, 2 cycles with a touchdown annealing temperature of 63°C and 62°C, respectively for 30 seconds, and 38 cycles with an annealing temperature at 61°C. Final DNA elongation was at 72°C for 10 minutes. DNA bands were visualized in prestained (0.4-μg/mL ethidium bromide) 3% agarose gels that were run for 1 hour at 120 V

### Statistical Analysis

Analyses were conducted using the statistical software Grand Prism 5 (San Diego, CA, USA). Means and SD were calculated for each studied parameter and an alpha level of 0.05 was selected throughout the study. The differences in psychometric dimensions across the three 5HTT-LPR genotype groups (l/l, l/s, s/s) were compared using ANOVA test (post-hoc: Bonferroni correction). Spearman’s correlations analysis were used to examine the relationship between personality dimensions, anxiety/depression symptoms, and coping subscales. Conformity of empirical genotype frequency distribution to theoretically expected Hardy–Weinberg equilibrium was verified using the Pearson χ2 test.

### Mediation analyses

A path method was applied to identify the potential mediation effects of maladaptive coping strategies and trait neuroticism on the relationship between 5-HTTLPR genotype and symptoms of anxiety and depression. The multiple-mediation model used is depicted as a path diagram in Figure 1. According to the general notations in the mediation analysis [80], we denote the total effect of the predictor on the outcome variable as *c*, which is the regression coefficient of the outcome on the predictor. We denote the effect of the predictor on the mediator as *a*, which is the regression coefficient of the mediator on the predictor. We denote the effect of the mediator on the outcome as *b*, which is the regression coefficient of the outcome on the mediator, and we denote the direct effect of the predictor on the outcome as *a’, b’, c’*, which is obtained by regressing the outcome on the predictor controlling for the mediator. Finally, we denote the estimated association among both mediators as *d*. We conducted the mediation analysis based on the procedure described by Baron & Kenny [80]. This approach involves testing multiple regression equations. First, the outcome variable is regressed on the predictor. Second, the mediator is regressed on the predictor variable. In the third equation, the outcome variable is regressed on both the predictor and the mediator. Mediation can be supposed to occur when the predictor (e.g., 5HTT-LPR genotype) significantly affects both the proposed mediator (e.g., maladaptive copying styles) and the outcome (i.e. symptoms of anxiety and depression) in the absence of the mediator, the mediator has a significant effect on the outcome, and the effect of the predictor on the outcome is reduced on the addition of the mediator to the model [81]. Specifically, we studied four different models: (i) the mediating effect of neuroticism on the relationship between the 5HTT-LPR genotype and symptoms of anxiety and depression; (ii) the mediating effect of maladaptive coping strategies on the relationship between the 5HTT-LPR genotype and symptoms of anxiety and depression; (iii) the mediating effect of maladaptive coping strategies on the relationship between neuroticism and symptoms of anxiety and depression and (iv) the mediating effect of neuroticism on the relationship between the 5HTT-LPR genotype and maladaptive coping strategies. The genetic variant was coded assuming an additive model (0 for ‘ll’, 1 for ‘ls’, and 2 for ‘ss’) as in our initial study [19]. Multicollinearity among predictors was assessed using a conventional tolerance threshold of <0.20. The significance of the indirect effects (mediating effect *ab*) is tested using the Partial Posterior (PP) method, which provides high statistical power while maintaining good Type I error control even under nonnormal data, coupled with a confidence interval (CI) obtained through a Hierarchical Bayesian approach [82, 83]. The joint test of significance was used to quantify the effect size of the indirect effects. It has been suggested that the value of the indirect effect relying on joint significance testing can be interpreted similar to squared correlation coefficients, where .01, .09, and .25 are considered small, medium, and large effect sizes [84].

## Results

The average age of the participants was 23.36 (SD = 8.48). Ages ranged from 18–36 years. Allele frequencies in the full samples did not differ significantly from those predicted by Hardy–Weinberg equilibrium (p > 0.05). A one-way ANOVA across the three 5HTT-LPR genotype groups (l/l, l/s, s/s) indicated a significant main effect of genotype on Neuroticism (F (2,130)=6.40, p=0.002). There was no significant association with the other sub-scales of the NEO-FFI. Post-hoc analyses showed an increase in neuroticism scores in the s/s genotype group (mean = 22.8, SD = 6.4) compared to the s/l (mean = 17.6, SD = 8.9)(p=0.02) and l/l (mean = 15.2, SD = 6.5) (p=0.001) groups. with regards to coping strategies, the ANOVA analysis revealed a main effect of genotype on focus on and venting of emotions (FVE) strategy (F (2,130)=4.90, p=0.008), with an increase in the FVE score in the s/s group (mean = 10.6, SD = 1.5) compared to s/l (mean = 8.9, SD = 2.5) (p=0.02) and l/l (mean = 8.6, SD = 2.7) (p=0.009) groups. There was no significant association with the other sub-scales of the COPE. With regards to symptoms of anxiety and depression the analyses indicated a main effect of genotype on Cognitive Anxiety (CA) (F(2,130)=8.14, p=0.0004) and on Emotional Arousal-Control (EAC) (F(2,130)=4.15, p=0.01). There was no significant association with other subscales of the IPPS-48 and the POMS. Post-hoc analyses revealed an increase in the CA score in the s/s group (mean = 22.0, SD = 6.6) compared to the l/l (mean = 15.0, SD = 5.1) (p=0.0003) group, but no significant difference between the s/s and the s/l (mean = 18.4, SD = 7.8) (p=0.08) groups. A post-hoc analysis showed a significant decrease in EAC in the s/s (mean = 21.1, SD = 2.2) compared to the l/l (mean = 24.4, SD = 3.8) (p=0.04) group, but no significant difference between the s/s and the s/l (mean = 22.0, SD = 5.6) (p=0.1). The results of the Spearman correlation analyses indicated that Neuroticism was significantly related to FVE (rho=0.42), CA (rho=0.50) and EAC (rho=0.38); FVE was significantly related to CA (rho=0.24) and EAC (rho=-0.32), (all p < 0.05).

### Mediation Analysis

All results from the mediation analysis are reported in Table 1. In the first model, the predictor variable is the 5HTT-LPR genotype. The outcome variables are symptoms of anxiety and the mediator variable is Neuroticism. The effect of the 5HTT-LPR-s genotype was statistically significant on the variables CA and EAC (path c). The effects of the 5HTT-LPR-s genotype on Neuroticism (path a^1^) and of Neuroticism on CA and EAC were also statistically significant (path b^1^). Therefore, the first three steps for mediation analysis, as described by Baron and Kenny, were satisfied. Our analysis based on the Partial Posterior method concluded that Neuroticism mediated the relationship between the 5HTT-LPR-s genotype and CA (p = 0.006; 95% CI = 0.046/0.198) with an effect size of 0.9. Neuroticism only partially (a^1^b^1^/(a^1^b^1^+c’) = 35% explained the effect of genotype on CA. The analysis revealed that Neuroticism also mediated the relationship between the 5HTT-LPR-s genotype and EAC (p = 0.001; 95% CI = -0.166/-0.031) with an effect size of 0.9. Neuroticism partially explained the effect of genotype on EAC (a^1^b^1^/(a^1^b^1^+c’) = 30%.

**Tab. 1.**
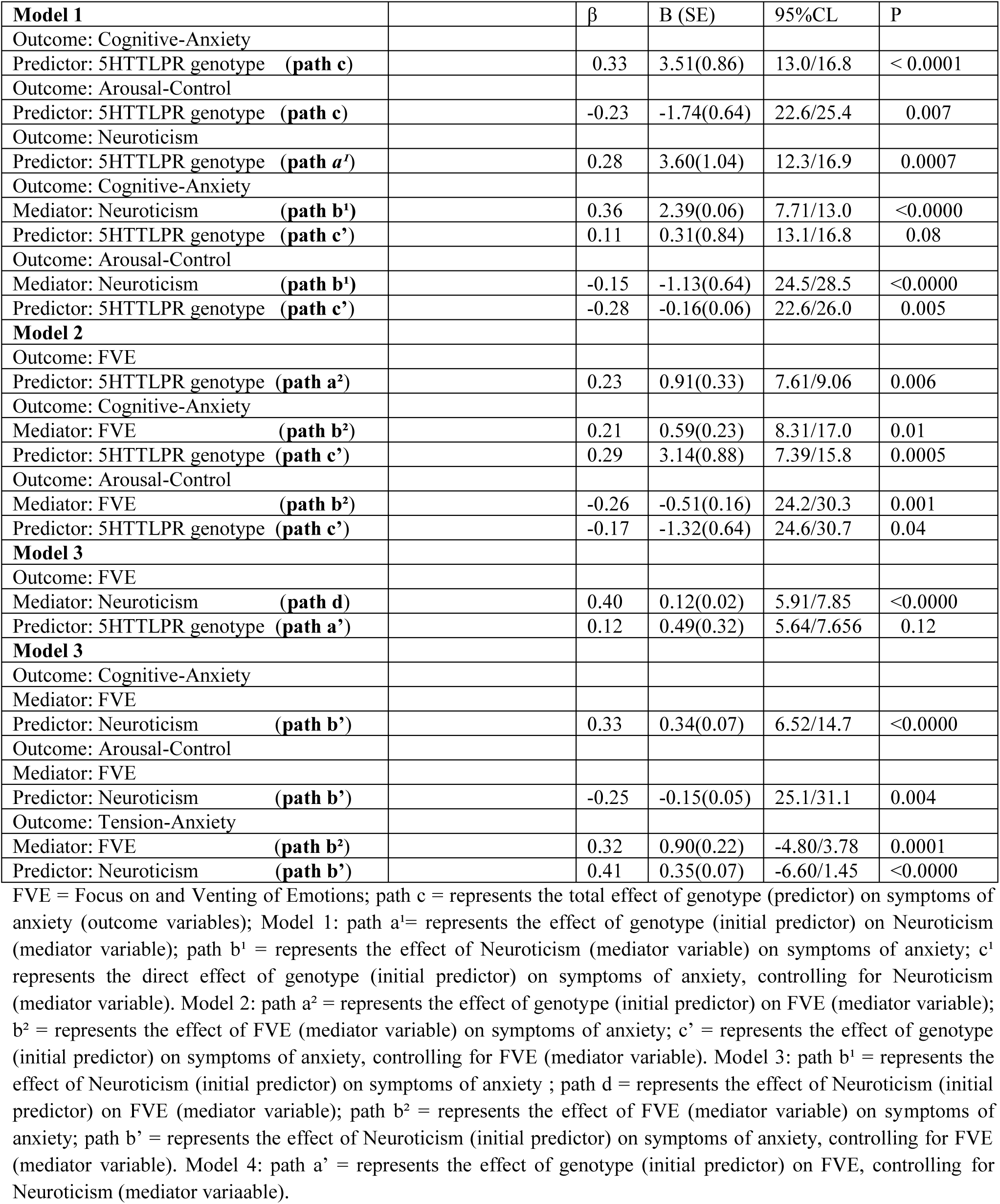
Results of Mediation Analysis

In the second model, we studied the effect of maladaptive coping (mediator) on the association between the 5HTT-LPR genotype and symptoms of anxiety. The effects of the 5HTT-LPR-s genotype on CA and EAC were the same as in the first model. We found a statistically significant effect of the 5HTT-LPR-s genotype on FVE (path a^2^) and of FVE on CA and EAC (path b^2^). The results of the Partial Posterior test showed that FVE mediated the relationship between the 5HTT-LPR-s genotype and CA (p = 0.01, 95% CI = 0.007/0.107) with an effect size of 0.5. FVE accounted for only (a^2^b^2^/(a^2^b^2^+c’) = 12% of the total effect of genotype on CA. FVE also mediated the relationship of the 5HTT-LPR-s genotype and EAC (p =0.006, 95% CI = -0.124/-0.013) with an effect size of 0.6 and accounted for (a^2^b^2^/(a^2^b^2^+c’) = 22% of the total the effect of genotype on EAC.

The third model tested the effect of maladaptive coping strategies on the relationship between Neuroticism and symptoms of anxiety. The effect of Neuroticism on T/A was significant. The effect of Neuroticism on the variables CA and EAC were the same as in the first model (path b^1^). The effect of Neuroticism on FVE (path d) and of FVE on the variables CA, EAC and T/A were significant (path b^2^). The application of the Partial Posterior test yielded the following results: FVE mediated the relationship between Neuroticism and T/A (p<0.0001, 95% CI=0.055/0.218) with an effect size of 0.9 and explained (db^2^/(db^2^+b’) = 22% of the Neuroticism effect on T/A; FVE was a mediator for Neuroticism and CA (p=0.01, 95% CI=0.019/0.164), with an effect size of 0.7, and explained (db^2^/(db^2^+b’) = 16% of the effect of Neuroticism on CA; FVE was a mediator for Neuroticism and EAC (p =0.006, % 0.95 CI = -0.124/-0.013) with an effect size of 0.6, and accounted for (db^2^/(db^2^+b’) = 28% of the effect of Neuroticism on EAC. Although Neuroticism and FVE were highly correlated, the associated tolerance of 0.83 suggests sufficient independence of these variables to enter them in the same model.

The aim of the last model in our study was to examine the indirect effect of Neuroticism on the relationship between the 5HTT-LPR genotype and maladaptive coping. In this situation, the independent and outcome variables were the 5HTT-LPR genotype and FVE, and the mediator was Neuroticism. The effect of the 5HTT-LPR-s genotype on FVE was the same as in the second model (path a^2^). The effects of the 5HTT-LPR-s genotype on Neuroticism (path a^1^) and of Neuroticism on FVE (path d) were the same as in the previous models. The mediation analysis revealed that Neuroticism mediated the relationship between the 5HTT-LPR-s genotype and FVE (*p* = 0.006; 95% CI = 0.046/0.198) with an effect size of 0.9. Neuroticism explained about one-third (a^1^d/(a^1^d+a’) = 32% of the effect of the 5HTT-LPR-s genotype on FVE.

## Discussion

Current evidence suggests that the short allele of the serotonin transporter promoter region (5-HTTLPR-s) appear to contribute to individual vulnerability to develop stress-induced psychopathologies such as symptoms of anxiety and depression through a variety of pathways. Particularly, direct or indirect effects mediated by anxiety related personality traits and negative coping strategies have been identified [39, 40]. In the present study, we performed a mediation analysis to investigate the mediating effect of Neuroticism and maladaptive coping strategies on the relationship between the 5HTT-LPR-s genotype and risk of developing symptoms of anxiety among elite athletes. To the best of our knowledge, this is the first report to explore the relationship between genetic factors, coping behavior and personality traits which influence the development of stress-related anxiety and depression symptoms in elite athletes. Four models were investigated.

In the first model we showed that the 5HTT-LPR s-allele is directly associated with symptoms of anxiety (i.e., increased cognitive-anxiety and low levels of arousal control), as well as with symptoms of anxiety through Neuroticism. These findings are consistent with the results of previous studies that reported a significant association between the 5HTT-LPR-s genotype and anxiety-related phenotypes including symptoms of anxiety and anxiety-related personality traits such as neuroticism [27, 29, 31, 85]. Recently, a number of studies suggested that 5-HTTLPR differences in anxiety and depression symptoms emerge significantly in the context of acute stress [33, 34]. Therefore, it is conceivable that athletes with one or two copies of the s allele are more likely then their counterparts with two copies of l allele to became anxious in response to stressful competitive experiences. Our data are also in line with findings that show how Neuroticism mediates the association between the 5HTTLPR-s genotype and affective disorders [39, 86] and support our previous investigation which suggested that the influence of the 5HTT-LPR s-allele on anxiety and emotion regulation among elite athletes is partially mediated by Neuroticism [19].

We further explored the effect of the 5HTT-LPR-s genotype on stress-related maladaptive coping strategies. Our findings suggest that the 5HTT-LPR-s allele is directly associated with FVE. Individuals with either one or two copies of the short 5HTT-LPR variant had significantly higher levels of FVE than those homozygous for the long genotype. Since no study to our knowledge has examined the influence of the 5HTT-LPR-s genotype on coping behavior in athletes, there is no empirical literature that directly informs the observed findings. Results, however, are in line with previous findings which suggest that the 5HTT-LPRs-allele could influence individual trait coping styles such as negative coping strategies (e.g., get angry, self-blame, crying alone) [40]. One possibility is that the 5HTT-LPR-s genotype exerts long-term effects on the development of neural structures central to emotional behavior as well as on adult brain plasticity and, thus, may influence broad classes of behavioral response to emotional stimuli, including individual coping strategies [61]. Several previous studies demonstrated that elite athletes who report more maladaptive coping strategies, experience greater symptoms of anxiety and depression [7, 59, 60]. We extended these original observations by providing evidence that emotion-focused maladaptive coping strategies also mediate the relationship between the 5HTT-LPR-s genotype and symptoms of anxiety. Our results also suggest that Neuroticism and FVE independently mediate the effect of the 5-HTT-LPR x stress interaction on anxiety symptoms. This is in line with results of Wang et al. [39] who showed that personality traits and negative trait coping styles might independently mediate the effect of the 5-HTT-LPR-s allele on affective disorders. However, interestingly, we found Neuroticism to be a stronger mediator than FVE in explaining the relationship between the 5HTT-LPR-s genotype and CA (p = 0.006 vs p = 0.01) and EAC (p = 0.001 vs p = 0.006).

Furthermore, our results suggest that Neuroticism is a mediating phenotype of the relationship between the 5-HTT-LPR-s genotype and maladaptive coping strategies. Importantly, we found that the 5-HTT-LPR-s genotype has a direct association with FVE, as well as an indirect association through its effect on neuroticism. This finding emphasizes the role and importance of Neuroticism and the way it influences coping strategy selection. Indeed, numerous previous studies have demonstrated that Neuroticism does influence coping strategy selection directly, by facilitating use of specific strategies, or indirectly, by influencing the frequency of exposure to stressors, the type of stressors experienced and the perception of stress [see for review, 67, 68]. Thus, if an association exists between the 5HTT-LPR-s genotype and maladaptive coping strategies, this may be the mechanism by which Neuroticism mediates the association between the 5HTT-LPR-s genotype and anxiety symptoms.

Finally, not surprisingly, results showed that FVE is also a potential mediating phenotype of the effect of Neuroticism on symptoms of anxiety. Previous studies have demonstrated that the use of maladaptive forms of emotion regulation mediated the association between Neuroticism and stress related outcomes (i.e., symptoms of depression) [69, 70, 87]. Our findings, thus, provide additional support for the notion that the use of maladaptive coping strategies might mediated the association between Neuroticism and emotional outcomes in a potentially stressful situation. These results are consistent with findings from studies reporting that a part of the influence of Neuroticism on stress related outcomes was mediated by maladaptive emotion focused coping [88]. It could be argued that these effects may be due to the multicollinearity between neuroticism and negative coping strategies. However, this is not the case in the current study, since multicollinearity statistics indicate independence of these variables. This is in line with prior evidence that coping explains variance in stress related outcomes in addition to that explained by personality [89].

Our study has several limitations worth noting. The first of these is that an unstructured interview was used to assess stressful events and therefore may be less accurate than structured stress interviews. Further research would benefit from using structured stress interviews. Second, the present study only examined the relationship between psychological raits and genetic influences on anxiety symptoms in male athletes. Thus, future research is necessary to test whether the findings in the present study can be generalized to women. Third, the non-clinical sample we used limits the generalizability of findings to athletes with anxiety disorders. Fourth, in this study, we did not find any association between the 5-HTT-LPR-s genotype and symptoms of depression. The relationship between the 5-HTT-LPR-s genotype and these aspects of negative emotionality was expected based on previous studies showing that the short allele was associated with higher depression scores [28, 90]. It is possible that the results of these previous studies be due to an interaction between stressful life events and 5-HTTLPR, whereby carriers of the s allele are more likely to suffer from depressive symptoms but only if they had experienced past traumatic events. Therefore, future studies need to take other adverse life events, potentially contributing to the incidence and severity of depressive symptoms, into consideration. Fifth, we analyzed one specific type of 5-HTTLPR allele variation. Several other single nucleotide polymorphisms in 5-HTT have been reported to be associated with symptom of depression [28, 91]. Therefore, future studies should examine other genetic variants and their interaction with psychological factors to influence anxiety/depression outcomes following acute stress in competitive sport. In conclusion, our results showed that the 5-HTT-LPR-s genotype has a direct effect on neuroticism, maladaptive coping and symptoms of anxiety and, most importantly, influences susceptibility to anxiety symptoms through neuroticism and focus on and venting of emotions. This can offer a new direction for future research to obtain more precise insights into these complex relationships and help further understand the complex nature of pre-competition anxiety among athletes. Finally, Neuroticism mediated the association between the 5-HTT-LPR-s genotype and focus on and venting of emotions and there also is also evidence for a role of focus on and venting of emotions in mediating the association between neuroticism and symptoms of anxiety. This knowledge is important when trying to determine possible intervention strategies for improving coping strategy in order to overcome sport related anxiety symptoms.

